# Within- and Between-Assessor Reliability of Lower-Limb Inter-Joint Coordination During Gait in Individuals With and Without Cerebral Palsy

**DOI:** 10.1101/2025.05.01.651648

**Authors:** Cloé Dussault-Picard, Yosra Cherni, Mickael Fonseca, Lena Carcreff, Fabien Leboeuf, Stéphane Armand

**Author notes:** **Corresponding author:** Cloé Dussault-Picard. **Declarations of interest:** none.

## Abstract

**Background:** Inter-joint coordination plays a key role in walking, particularly in people with cerebral palsy (CP), who experience altered movement patterns. The Continuous Relative Phase (CRP) method quantifies lower-limb coordination by assessing the phase relationships between joints. However, the reliability of CRP measurements during walking in individuals with CP remains unexplored, and may be affected by measurement variability due to marker placement errors, soft tissue artifacts, and natural movement fluctuations. Quantifying this reliability is important for appropriate clinical comparisons. This study aimed to quantify within- and between-assessor reliability of lower-limb CRP measurements in individuals with CP and their non-impaired (NI) peers.

**Methods:** CP (n=19, age=18.4±7.3 years, GMFCS I–III) and NI (n=19, age=18.3±11.2 years) individuals completed two gait assessment sessions, each including 3D motion capture of at least 10 walking trials. Two trained assessors independently placed reflective markers and conducted gait analyses. Standard error of measurement (SEM) and minimal detectable change (MDC) were computed for knee-hip and ankle-knee coordination across gait subphases.

**Findings:** The SEM and MDC were lower for knee-hip than ankle-knee coordination, suggesting higher measurement reliability for proximal joint coupling. For knee-hip coordination, MDC reached 15.1±0.7°(CP) and 9.3±0.6°(NI) between assessors, and 23.8±3.0°(CP) and 9.1±1.6°(NI) within assessors. For ankle-knee coordination, MDC reached 29.0±2.6°(CP) and 25.0±3.5°(NI) between assessors, peaking at 47.3±10.9°(CP) and 28.6±1.5°(NI) in mid-swing within assessors.

**Interpretation:** This study provides the first metrological reference for reliability of CRP-based inter-joint coordination during gait in CP. Results showed poor reliability, emphazing that such measurements must be interpreted with caution.

**Highlights:**

- Knee-hip showed greater reliability than ankle-knee coordination across gait phases
- Cerebral palsy individuals showed higher variability than non-impaired peers

The beginning and the end of gait cycle showed the poorest reliability

Pre-post comparisons should account for MDC thresholds to avoid misinterpretation

## 1. Introduction

Cerebral palsy (CP) refers to neuromotor disorders that occur in approximately 2 births per 1000 children (Oskoui et al., 2013). CP is characterized by progressive neuromusculoskeletal impairments, such as bony torsions and contractures, that contribute to walking deviations (Armand et al., 2016), limited participation (Gjesdal et al., 2020) and reduced functional independence (Elad et al., 2018; Schmidt et al., 2020) compared to non-impaired (NI) peers.

Three-dimensional gait analysis contributes to identifying walking impairments of individuals with CP (Baker, 2006). By estimating gait parameters such as kinematics and kinetics, these analyses provide a comprehensive understanding of the mechanics of movement. However, walking is a complex activity that relies on the continuous coordination of multiple joints (E.A van Emmerik et al., 2014). As a result, there has been a growing interest over the past decade in exploring higher-order metrics that extend beyond traditional joint kinematics such as inter-joint coordination (Carollo et al., 2018; Chiu et al., 2015; Chiu and Chou, 2012; Dussault-Picard et al., 2023, 2022; Fowler et al., 2010; Ippersiel et al., 2021a, 2021b), which may reveal deeper insights into movement patterns and their impairments. Inter-joint coordination provides valuable information on how joints interact to produce a coordinated and efficient movement (Byrne et al., 2002; E.A van Emmerik et al., 2014) to better evaluate motor strategies and compensatory adaptations in individuals with CP during walking (Dussault-Picard et al., 2023, 2022).

Based on dynamical systems theory, movement coordination can be represented by the phase space of the system, where each state is described by a time-dependent signal (e.g., a segment or joint angle) and its first derivative, representing velocity. The phase space allows for identifying key features of the system’s behavior, such as its stability and ability to change, and potential for transitions or bifurcations under changing environmental or task constraints (E.A van Emmerik et al., 2014). Lamb and Stöckl (2014) have proposed a method for representing the phase space of gait kinematics signals by accounting for the amplitude and frequency differences (Lamb and Stöckl, 2014). This method relies on centering the amplitude of the data around zero (Rosenblum et al., 2001) and transforming the signal using a Hilbert transformation rather than using its first derivative (i.e., angular velocity) (Lamb and Stöckl, 2014).

Inter-joint coordination can be quantified by the phase difference between two joint kinematic signals using the continuous relative phase (CRP) (Burgess-Limerick et al., 1993). This metric represents the in-phase/out-of-phase coupling relationships, where a value of 0° refers to two joints that are moving fully in phase with each other, whereas a value of 180° indicates a fully out-of-phase coupling (E.A van Emmerik et al., 2014). It has been shown that individuals with CP exhibit more in-phase motion than NI peers (Carollo et al., 2018; Dussault-Picard et al., 2023, 2022; Fowler et al., 2010), and when walking on uneven surfaces (Dussault-Picard et al., 2023), which was associated with a more restricted and rigid walking pattern. Despite consistent and significant differences in coordination between groups and/or conditions, the clinical relevance of these differences remains uncertain (Paiva et al., 2024), given the current lack of evidence supporting the reliability of this measurement approach. Variability in inter-joint coordination may arise from marker placement, soft tissue artifacts, natural fluctuations in movement patterns, or individual adaptations (e.g., fatigue). Quantifying the reliability of measurements across different sessions and assessors is essential for accurately interpreting and contextualizing the clinical significance of group and/or condition differences.

To our knowledge, the reliability of inter-joint coordination during walking has yet to be investigated, particularly in individuals with CP. Thus, this study aimed to quantify within-assessor and between-assessor standard error of measurement (SEM), and the minimal detectable change (MDC) values of the lower-limb inter-joint coordination using the CRP method in individuals with CP and their NI peers.

## 2. Methods

### 2.1. Participants

Participants with and without CP were recruited at the Geneva University Hospitals (aged between 6-43 years old). Individuals with CP were included if they had a confirmed diagnosis of CP, the ability to walk 10 meters without assistive aid, and the capacity to understand and follow verbal instructions. The NI individuals were included if they had no known pathological condition affecting typical motor ability. Individuals were excluded if they had a known pregnancy and allergy to adhesive tape. The research protocol was approved by the “Commission Cantonale d’Éthique de la Recherche Genève” (CCER-2020-00358) and all the participants or their legal tutors for non-adult participants provided written informed consent.

### 2.2. Experimentation

Participants attended the laboratory on two separate assessment sessions (S1 and S2), spaced 10 days apart. During the first session (S1), a 3D gait analysis session was performed by a first assessor (A1) (S1A1). The second visit session (S2) involved two consecutive 3D gait analysis assessments, conducted by the first (A1) and a second (A2) assessors, labeled as S2A1 and S2A2, respectively. Each 3D gait analysis assessment included at least 10 barefoot walking trials at self-selected comfortable pace without assistive aid. The assessor conducting the session was responsible for placing the reflective markers (14mm), following the Van Sint Jan (2007) guidelines (Sint Jan, 2007) and according to the Conventional Gait 2 Model (version 2.3) (Leboeuf et al., 2019). Both assessors were highly experienced in locating anatomical landmarks. The marker trajectories were recorded at 100Hz by a 12-camera motion capture system (Oqus7+, Qualisys, Go□teborg, Sweden).

### 2.3. Data processing

Initial data processing (i.e., marker labelling and gap filling, foot strike and foot-off detection) was conducted in QTM software version 2022.3 (Qualisys, Go□teborg, Sweden). The Zeni algorithm was used for foot strike and foot off identifications (Zeni et al., 2008), and manual verification was carried out. The hip, knee and ankle joint kinematics were calculated using the CGM2.3 lower-limb version of the open source library pyCGM2 (Leboeuf et al., 2019). The c3d files were then exported and further processed in MATLAB (The MathWorks, Inc, vR2023b) using the BiomechZoo Toolbox (Dixon et al., 2017). The CRP was calculated for the knee-hip (KH) and ankle-knee (AK) coupling in the sagittal plane. Phase angles of each joint were calculated following the Lamb and Stöckl (2014) recommendations (Lamb and Stöckl, 2014):

4) First, the range of the signal’s x(*t*) amplitude was centered around zero (Eq. 1):

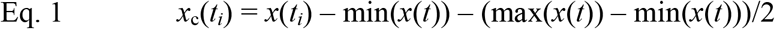
2) Based on the centered signal *x*_*c*_(*t*), an analytic signal z(*t*) was created using the Hilbert transform, where *H*(*t*) of *x*_*c*_(*t*) is the imaginary part of the analytic signal (Eq. 2):

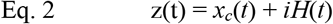
3) The phase angle □ at time *t*_*i*_ was then calculated (Eq. 3):

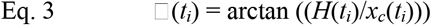
4) Afterward, the CRP analysis was used to describe inter-joint coordination by calculating the absolute difference in the phase angle of the distal joint □ _*2*_ from that of the proximal joint □ _*1*_ across the movement cycle (Eq. 4):

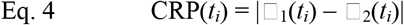

Then, the CRP curves were time-normalized to 101 data points according to foot contacts. For individuals with unilateral CP, the most affected leg was kept for analysis whereas both legs were considered separately for bilateral CP and NI individuals.

### 2.4. Statistical analysis

The frame-by-frame (*i*) standard error of measurement (SEM) was calculated for each metric and group (KH-CP, KH-NI, AK-CP, AK-NI):

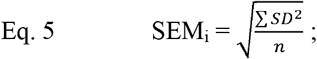

where SD is the standard deviation of assessment averages at frame (*i*) and *n* is the number of measurements within each assessment (i.e., number of participants).

The MDC for between assessor and within assessor measurements was calculated for each metric as:

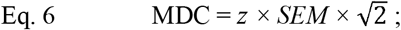

where z-score (*z*) corresponds to 1.96 and *SEM* is the SEM curve (between assessors or within assessors). The MDC curves were then averaged across each gait subphase (Baker, 2013): first double support, early single support, mid-single support, late single support, second double support, early swing, mid-swing, late swing. All statistical analysis steps were run in Rstudio (v.2024.04.2, PBC, Boston, MA).

## 3. Results

### 3.1. Participants

A total of 19 individuals with CP (Gross Motor Function Classification System (GMFCS) I: n = 15, GMFCS II: n = 3, GMFCS III: n = 1) and 19 NI individuals were recruited. Anthropometrics and characteristics of each group are reported in **Table 1**.

**Table 1.** Participant characteristics.

|  | CP | NI |
| --- | --- | --- |
| n | 19 | 19 |
| Age | 18.4 (7.3) | 18.3 (11.2) |
| Sex | 5F / 14M | 9F / 10M |
| BMI | 21.0 (4.2) | 19.3 (3.7) |
| Height (m) | 1.6 (0.2) | 1.5 (0.2) |
| Weight (kg) | 56.1 (17.9) | 46.9 (18.1) |
| GMFCS level | I: n = 15 |  |
|  | II: n = 3 | NA |
|  | III: n = 1 |  |
| Involvement | Unilateral: n = 9 | NA |
|  | Bilateral: n = 10 |  |

Mean (standard deviation) of participant anthropometrics and characteristics for the cerebral palsy (CP) and the non-impaired (NI) groups. Sample size (n), side of the muscle disorder involvement; unilateral (U) or bilateral (B), Gross Motor Function Classification System (GMFCS), not applicable (NA), and sex; female (F) and male (M) are also reported.

### 3.2. Knee-hip coordination reliability

The between-assessor SEM of the KH coordination is greater at the beginning of the gait cycle, namely during the first double support (MDC-CP: 15.1 ± 0.7°; MDC-NI: 9.3 ± 0.6°) and early single support (MDC-CP: 10.4 ± 2.5°; MDC-NI 8.5 ± 2.6°) (**Figure 1**). This results in an average maximal difference between the CRP curves of 8.4 ± 6.2° and 4.9 ± 3.1° during the first double support, and of 7.2 ± 4.6° and 6.6 ± 3.7° during the early single support for the CP and the NI group, respectively (**Table 2**).

**Table 2.** Within and between assessor differences between the continuous relative phase curves.

| Subphase | Event | Between assessors CRP difference |  |  |  | Within assessors CRP difference |  |  |  |
| --- | --- | --- | --- | --- | --- | --- | --- | --- | --- |
|  |  | KH CP | KH NI | AK CP | AK NI | KH CP | KH NI | AK CP | AK NI |
| **1DS** | **max** | 8.4 (6.2) | 4.9 (3.1) | 13.9 (13.0) | 15.0 (9.1) | 13.5 (11.6) | 5.4 (2.5) | 20.0 (21.4) | 21.2 (20.6) |
| **1DS** | **min** | 3.5 (3.0) | 2.1 (2.9) | 6.2 (9.9) | 4.8 (7.6) | 4.8 (5.6) | 2.0 (2.0) | 9.0 (12.8) | 8.3 (14.5) |
| **1DS** | **mean** | 5.9 (4.5) | 3.6 (3.0) | 9.6 (11.1) | 9.4 (8.1) | 8.7 (6.9) | 3.8 (2.3) | 13.9 (15.2) | 14.7 (17.3) |
| **ESS** | **max** | 7.2 (4.6) | 6.6 (3.7) | 13.2 (9.8) | 10.0 (7.2) | 12.3 (11.4) | 9.0 (6.7) | 17.6 (14.8) | 14.5 (12.4) |
| **ESS** | **min** | 1.6 (2.3) | 1.2 (1.2) | 5.1 (8.5) | 1.9 (3.2) | 4.0 (6.5) | 1.6 (1.8) | 4.6 (8.5) | 2.8 (6.8) |
| **ESS** | **mean** | 4.1 (3.1) | 3.7 (1.9) | 9.5 (9.3) | 5.9 (4.1) | 7.7 (8.1) | 5.3 (4.1) | 11.1 (10.3) | 8.9 (9.6) |
| **MSS** | **max** | 4.5 (2.8) | 2.7 (1.6) | 12.9 (10.2) | 5.5 (3.6) | 6.7 (6.0) | 4.0 (3.9) | 11.5 (9.2) | 7.0 (6.8) |
| **MSS** | **min** | 2.1 (2.1) | 0.9 (1.0) | 5.0 (6.0) | 3.1 (2.6) | 2.8 (2.9) | 1.4 (2.4) | 4.6 (6.4) | 2.5 (3.4) |
| **MSS** | **mean** | 3.4 (2.3) | 1.8 (1.3) | 9.1 (7.5) | 4.3 (3.0) | 4.8 (4.4) | 2.5 (2.9) | 8.0 (7.7) | 4.7 (4.6) |
| **TSS** | **max** | 4.0 (2.6) | 2.4 (1.4) | 9.2 (6.8) | 3.8 (2.4) | 4.5 (3.1) | 2.5 (2.0) | 8.1 (6.4) | 4.3 (3.3) |
| **TSS** | **min** | 1.6 (2.2) | 0.7 (0.9) | 3.0 (3.3) | 1.8 (1.8) | 1.7 (1.9) | 0.8 (0.9) | 2.8 (3.2) | 1.7 (2.0) |
| **TSS** | **mean** | 2.8 (2.3) | 1.6 (1.1) | 5.9 (4.3) | 2.7 (1.9) | 3.0 (2.4) | 1.7 (1.3) | 5.3 (4.3) | 3.0 (2.4) |
| **2DS** | **max** | 3.4 (2.) | 2.5 (1.3) | 5.5 (3.4) | 3.9 (2.4) | 3.6 (2.6) | 2.1 (1.3) | 6.4 (5.8) | 4.6 (4.6) |
| **2DS** | **min** | 1.8 (2.3) | 1.5 (1.2) | 1.4 (2.1) | 0.7 (1.2) | 1.6 (1.9) | 1.1 (1.2) | 2.4 (4.5) | 0.6 (0.7) |
| **2DS** | **mean** | 2.6 (2.4) | 2.1 (1.2) | 3.6 (2.7) | 2.6 (1.8) | 2.6 (2.2) | 1.6 (1.2) | 4.8 (5.3) | 2.6 (2.2) |
| **ESw** | **max** | 3.9 (2.9) | 2.9 (1.6) | 7.7 (4.5) | 8.1 (4.2) | 4.7 (2.8) | 2.5 (1.4) | 11.1 (8.5) | 10.5 (11.1) |
| **ESw** | **min** | 2.3 (2.3) | 1.7 (1.7) | 1.4 (1.5) | 0.9 (2.2) | 2.3 (2.0) | 1.3 (1.2) | 2.0 (2.2) | 2.0 (3.1) |
| **ESw** | **mean** | 3.2 (2.7) | 2.3 (1.6) | 4.5 (2.3) | 4.2 (2.3) | 3.6 (2.4) | 2.0 (1.3) | 6.0 (4.3) | 6.3 (6.5) |
| **MSw** | **max** | 4.3 (2.6) | 3.5 (2.0) | 18.8 (9.5) | 11.5 (6.8) | 5.3 (2.6) | 2.6 (1.5) | 26.5 (25.9) | 14.0 (14.3) |
| **MSw** | **min** | 1.7 (2.2) | 2.0 (2.1) | 2.7 (4.2) | 3.6 (3.1) | 2.2 (2.6) | 1.2 (1.5) | 4.3 (5.3) | 5.0 (7.2) |
| **MSw** | **mean** | 3.0 (2.3) | 2.8 (2.0) | 10.2 (6.1) | 8.1 (5.0) | 3.7 (2.6) | 1.9 (1.5) | 16.0 (16.0) | 9.6 (10.4) |
| **LSw** | **max** | 5.5 (3.7) | 4.4 (2.5) | 20.9 (16.7) | 13.5 (9.4) | 8.5 (6.4) | 3.5 (1.7) | 24.4 (22.2) | 18.3 (22.7) |
| **LSw** | **min** | 1.9 (2.0) | 1.9 (2.0) | 5.9 (7.5) | 2.5 (2.7) | 3.7 (3.2) | 1.1 (1.3) | 4.6 (5.8) | 6.5 (12.0) |
| **LSw** | **mean** | 3.6 (2.5) | 3.1 (1.9) | 13.7 (13.4) | 7.9 (5.3) | 5.8 (4.0) | 2.3 (1.4) | 15.2 (15.0) | 12.0 (17.3) |

Group means (SD) of the continuous relative phase curves difference (in degrees) for the knee-hip (KH) and ankle-knee (AK) joint pairs during each subphase of the gait cycle: first double support (1DS), early single support (ESS), mid-single support (MSS), late single support (LSS), second double support (2DS), early swing (ESw), mid-swing (MSw), late swing (LSw). Maximal (max), minimal (min) and average (mean) differences for each metrics are presented for each group: cerebral palsy (CP) and non-impaired (NI) individual.

**Figure 1.**
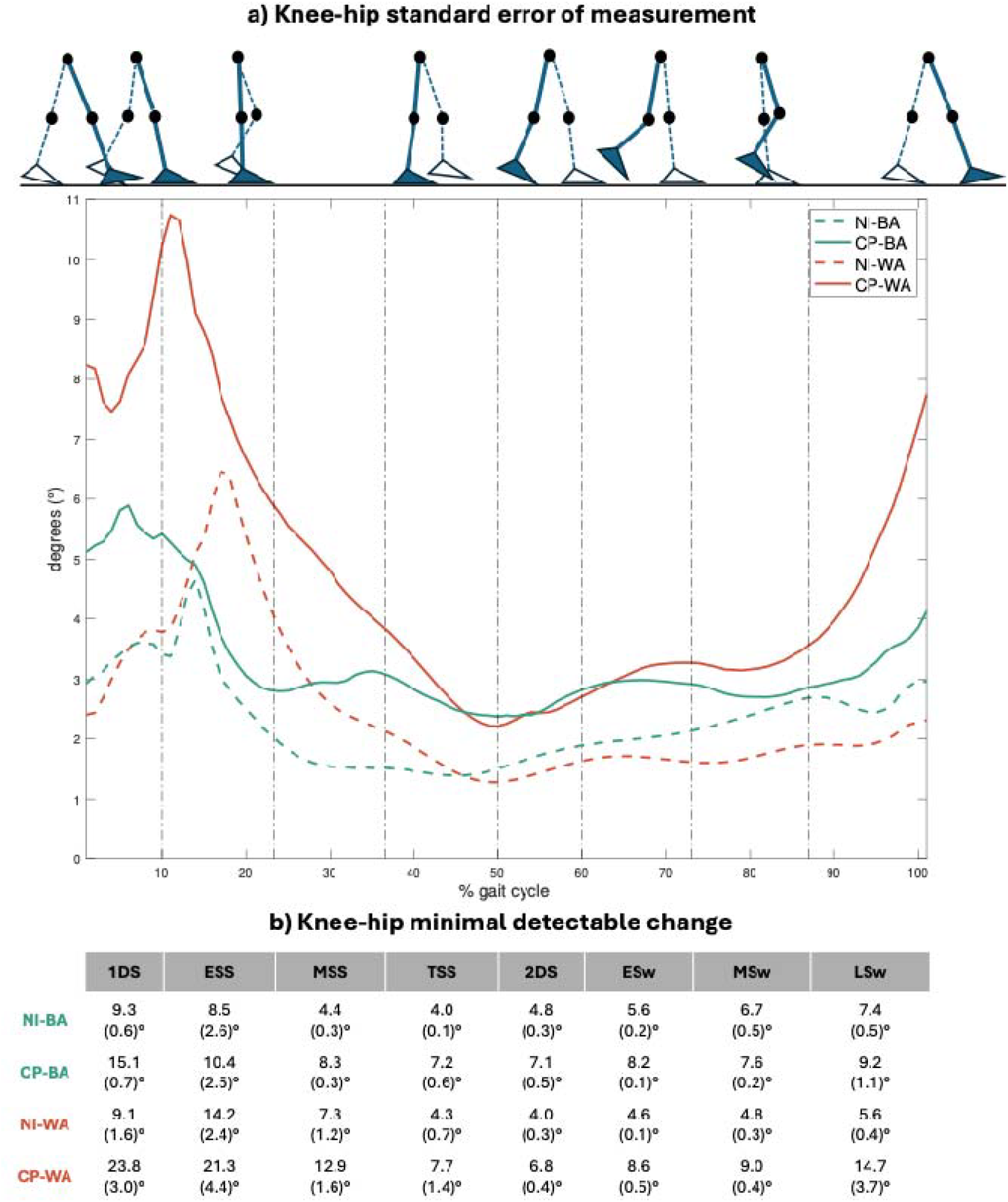
**a)** Continuous Standard Error of Measurement (SEM) of knee-hip coordination in cerebral palsy (CP) individuals (n = 29 legs) and non-impaired (NI) individuals (n = 19 legs) for within-assessor (WA) and between-assessor (BA) measurements. Each vertical dashed line delimited the gait cycle subphases, as described Baker (2013) (Baker, 2013); **b)** Mean (standard deviation) minimal detectable change (MDC) of knee-hip coordination during subphases of the gait cycle in both groups. Abbreviations: 1SD, first double support; ESS, early single support; MSS, mid-single support; LSS, late single support; 2SD, second double support; ESw, early swing; MSw, mid-swing; LSw, late swing.

The within-assessor SEM of the KH coordination is also greater during the first double support (MDC-CP: 23.8 ± 3.0°; MDC-NI: 9.1 ± 1.6°) and early single support (MDC-CP: 21.3 ± 4.4°; MDC-NI 14.2 ± 2.4°) (**Figure 1**). This results in an average maximal difference between the CRP curves of 13.5 ± 11.6° and 5.4 ± 2.5° during the first double support, and of 12.3 ± 11.4° and 9.0 ± 6.7° during the early single support for the CP and the NI group, respectively (**Table 2**).

### 3.3. Ankle-knee coordination reliability

The between-assessor SEM of the AK coordination is greater at the beginning and the end of the gait cycle, namely during the first double support (MDC-CP: 29.0 ± 2.6°; MDC-NI: 25.0 ± 3.5°), the mid-swing (MDC-CP: 25.7 ± 4.2°; MDC-NI: 19.4 ± 2.4°), and late swing (MDC-CP: 38.9 ± 3.0°; MDC-NI: 20.5 ± 6.0°) (**Figure 2**). This results in an average maximal difference between the CRP of 13.9 ± 13.0° and 15.0 ± 9.1° during the first double support, of 18.8 ± 9.5° and 11.5 ± 6.8° during the mid-swing, and of 20.9 ± 16.7° and 16.5 ± 9.4° during the late swing for the CP and the NI group, respectively (**Table 2**).

**Figure 2.**
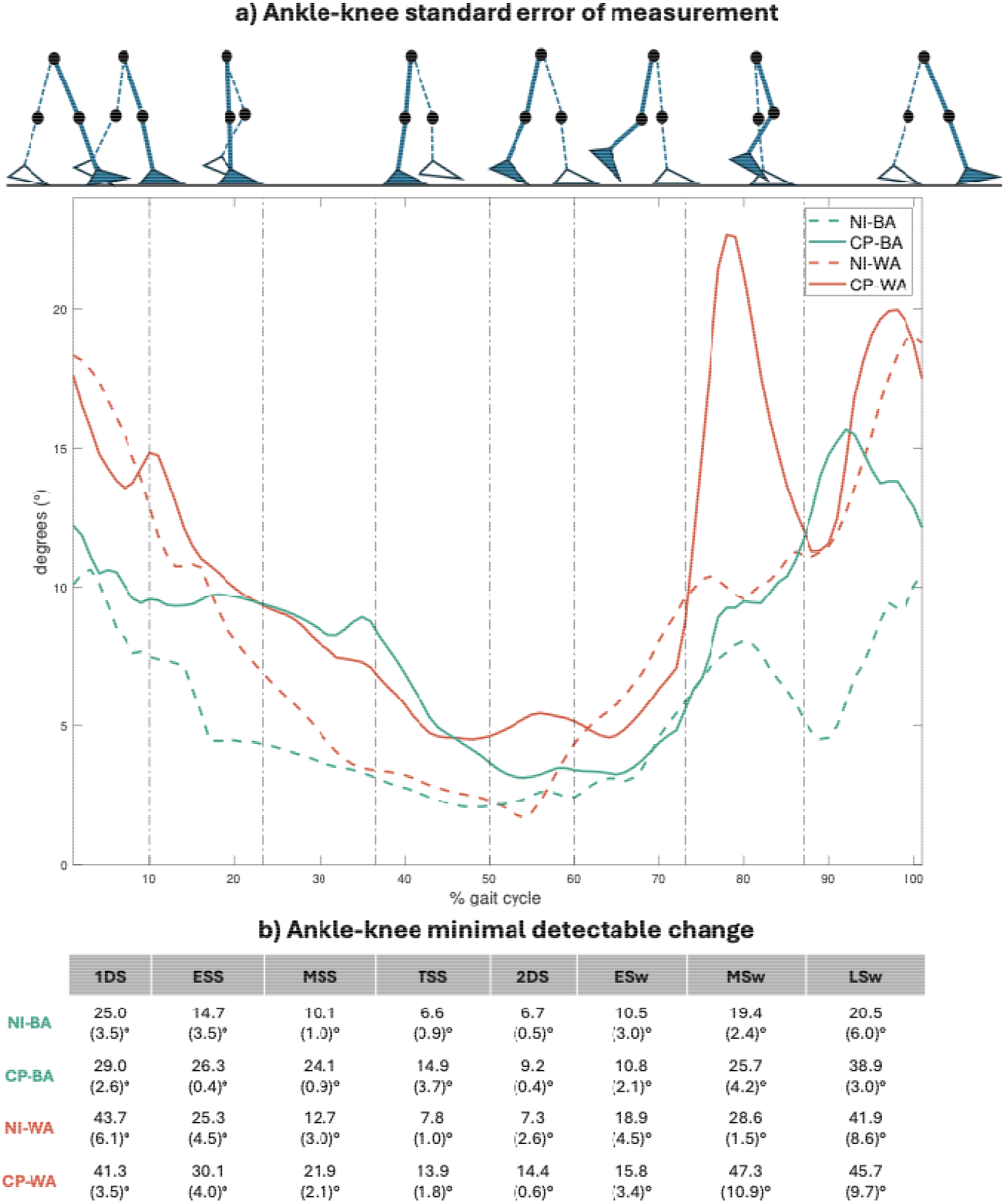
**a)** Continuous Standard Error of Measurement (SEM) of ankle-knee coordination in in cerebral palsy (CP) individuals (n = 29 legs) and non-impaired (NI) individuals (n = 19 legs) for within-assessor (WA) and between-assessor (BA) measurements. Each vertical dashed line delimited the gait cycle subphases, as described by Baker (2013) (Baker, 2013); **b)** Mean (standard deviation) minimal detectable change (MDC) of ankle-knee coordination during subphases of the gait cycle in both groups. Abbreviations: 1SD, first double support; ESS, early single support; MSS, mid-single support; LSS, late single support; 2SD, second double support; ESw, early swing; MSw, mid-swing; LSw, late swing.

The within-assessor SEM of the AK coordination is also greater during the first double support (MDC-CP: 41.3 ± 3.5°; MDC-NI: 43.7 ± 6.1°), the mid-swing (MDC-CP: 47.3 ± 10.9°; MDC-NI: 28.6 ± 1.5°), and late swing (MDC-CP: 45.7 ± 9.7°; MDC-NI: 41.9 ± 8.6°) (**Figure 2**). This results in an average maximal difference between the CRP curves of 20.0 ± 21.4° and 21.2 ± 20.6° during the first double support, of 26.5 ± 25.9° and 14.0 ± 14.3° during the mid-swing, and of 24.4 ± 22.2° and 18.3 ± 22.7° during the late swing for the CP and the NI group, respectively (**Table 2**).

## 4. Discussion

This study aimed to establish SEM and MDC values for within-assessor and between-assessor reliability of lower-limb inter-joint coordination using the CRP method in individuals with CP and their NI peers. Three main findings emerged. First, the SEM and MDC were lower for KH compared to AK coordination in both groups. Second, higher SEM and MCD were observed in patients with CP compared to NI individuals. Third, higher SEM and MDC values were observed during the first double support and early single stance for KH coordination, and during mid-swing for AK coordination.

### 4.1. Clinical interpretation

The present findings highlight the importance of considering measurement variability when interpreting pre-post changes in joint coordination using CRP analysis. The MDC reported in this study provides essential reference values to distinguish actual coordination modifications from measurement variability. When assessing improvements or deteriorations in inter-joint coordination following interventions or comparing data between groups, results should be interpreted with caution. For instance, MDC of the KH coordination (% of group mean amplitude during subphase) can reach more than 23.8° (96.9%) in CP and 9.3° (104.5%) in NI individuals during the first double support, while the MDC of the AK coordination can reach more than 41.3° (96.2%) in CP and 43.7° (118.3%) in NI individuals during the same subphase. The first double support subphase, however, was characterized by high inter-individual variability (Dussault-Picard et al., 2023), which may contribute to the difficulty in detecting statistically significant differences when comparing conditions or groups. As a result, even when observed changes approach or exceed the MDC thresholds, they may not always reach statistical significance because of the data distribution, highlighting the need for cautious interpretation.

### 4.2. Measurement procedures variability

While experienced assessors were involved in this study, the differences observed between their measurements highlight the potential variability induced by measurement procedures such as marker placement and gait event detection. This is particularly relevant for the CP group, where anatomical and functional deviations may exacerbate such effects (Everaert et al., 2023). Moreover, the choice of the kinematic model can also influence measurement variability, as different models may be sensitive to soft tissue artifact (Cockcroft et al., 2016; Leboeuf et al., 2023; Leboeuf and Sangeux, 2023). In the current study, the CGM2.3 lower-limb model was used (Leboeuf et al., 2019). This model optimized motion reconstruction algorithms by using kinematic fitting, which minimizes errors caused by soft tissue movement (Leboeuf et al., 2023). The numerous methodological variations across studies—including the coordinated elements (e.g., joints, segments), the chosen coordination analysis method (e.g., CRP, vector coding, power spectral density), and the movement performed (e.g., running, walking, jumping)—make it difficult to compare our results with previously published data. However, previous investigations of inter-assessor SEM for sagittal inter-segmental coordination during running, using the vector coding method in healthy adults, revealed that the maximum error occurred during mid-stance, with an average MDC of 15.3° for foot-tibia phase angles (Paiva et al., 2024).

### 4.3. Sensitivity of continuous relative phase analyses

Another key factor lies in the sensitivity of CRP analyses. Indeed, the **figure 3** shows that even minimal differences in kinematics (amplitude and/or timing) between two sessions conducted by the same assessor can lead to a consequent shift in the resulting CRP curve. This difference may be attributed to the sensitivity of the signal at points of bifurcation (or phase transitions), such as during the switch from flexion to extension, where angular velocity is low. These changes in coordination, also referred to as bifurcations or phase transitions, are particularly sensitive points where the stability of the system is challenged (i.e., the system is more sensitive and less stable) (E.A van Emmerik et al., 2014). In individuals with CP, reduced amplitude and limited motion at the ankle (de Bruin et al., 2013) further amplify these effects, making coordination patterns more variable and prone to measurement error (see **figure 3**). Indeed, high SEM for the AK coordination has been observed during the mid-swing subphase, reaching up to a MDC of 47.3° in CP individuals (**figure 2, figure 3**). As a result, studies on inter-joint coordination using the CRP method should consider that this subphase involves relatively high error, and the interpretation of significant results at these instants should be approached with caution. This high sensitivity also raises questions about the suitability of CRP analysis for assessing coordination in pathological populations such as individuals with CP, who often exhibit restricted ranges of motion (de Bruin et al., 2013) and temporal variability in gait (Dussault-Picard et al., 2025). Future studies should further investigate whether alternative methods of inter-joint coordination calculations (i.e., vector coding, power spectral density) and signal alignment (i.e., dynamic time warping) may provide more repeatable measures of coordination in populations with such characteristics. Indeed, it is common for certain physiological signal analysis methods to be better suited for specific populations, as demonstrated in the case of muscle synergy analysis (Rabbi et al., 2020).

**Figure 3.**
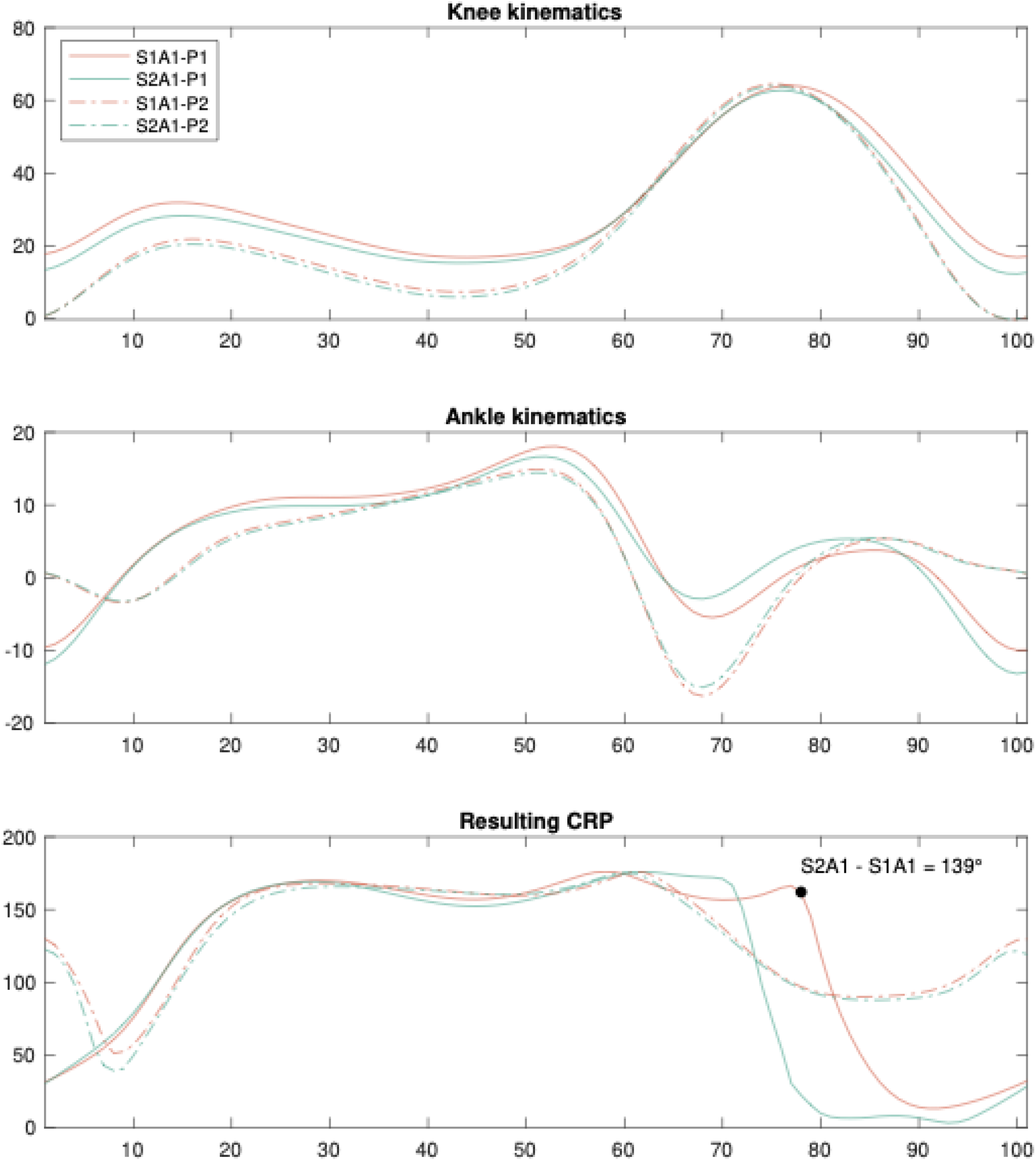
Example of two sessions (S1, S2) conducted by the same assessor (A1) for two participants (P1, P2), representing the effect of a temporal shift between kinematics of different sessions on the resulting continuous relative phase values. The P1 have cerebral palsy whereas P2 is unimpaired. The continuous relative phase difference between sessions reached up to 139° at 78% of the gait cycle for P1.

### 4.4. Cerebral palsy vs. non-impaired individuals

The CP group exhibited higher SEM and MDC values across both metrics, indicating greater variability and reduced repeatability in their inter-joint coordination patterns (Dussault-Picard et al., 2022). This variability may reflect the neuromotor impairments associated with CP, such as spasticity, altered voluntary control of movement, and compensatory strategies, which can lead to inconsistent joint coupling during gait (Dussault-Picard et al., 2022). The observed differences between groups highlight the need to investigate the factors underlying these discrepancies, such as functional level, age, or other individual characteristics.

### 4.5. Proximal vs. distal joints

In both groups, the SEM and MDC values were lower for KH compared to AK coordination, suggesting that KH coordination is more stable and reliable as a metric for assessing inter-joint coordination during gait. This result can be explained by two key factors: the range of motion and the number of directional changes in joint angles. The ankle undergoes more frequent directional changes during gait, as it constantly adjusts to ground contact and propulsion forces (Dumas and Cheze, 2008; Konor et al., 2012). These rapid and complex adjustments introduce greater variability in AK coordination, making it more sensitive to within- and between-assessor differences. In contrast, KH coordination involves fewer directional changes and follows a smoother, more predictable movement pattern, which likely contributes to its lower SEM and MDC values.Consequently, KH coordination appears to be a more stable and reliable outcome measure for tracking gait changes over time.

### 4.6. Study limitations

Several limitations should be acknowledged. First, the study included only 19 individuals with CP, most of whom have low motor impairments (GMFCS level I), which may limit the generalizability of the results. Second, the age range of participants was relatively wide (6–43 years), which introduces another potential source of variability, as age-specific differences in motor control and development could influence coordination (Seidler et al., 2010). Despite these limitations, our results provide baseline values, and further research should expand the metrological assessment of the CRP outcome to better understand measurement errors within the population, considering factors such as functional level and age.

## 5. Conclusion

This study provides an initial metrological reference for the reliability of the CRP during gait in individuals with and without CP. The results underscore the importance of standardized protocols and assessor training in minimizing kinematics variability, which may result in high SEM of the resulting CRP curves during specific subphases of the gait cycle (i.e. during the first double support and early single stance for KH coordination, and during the mid-swing for AK coordination). Future studies using CRP analysis to assess 12esponsiveness (e.g., pre-post intervention) or to compare groups should interpret results with caution and consider that a MDC greater than 20° may be relevant at the beginning of the cycle for hip-knee coordination, while a MDC greater than 40° may be considered at both the beginning and end of the cycle for AK coordination.

## Acknowledgements

We would like to acknowledge the Fonds de recherche du Québec —Nature et technologie for the postdoctoral scholarship of Cloé Dussault-Picard.

## Author contributions

Conceptualization; SA, CDP – Data acquisition; LC, MF – Formal analysis; CDP – Supervision; SA – Roles/Writing – original draft; YC, CDP – Writing – review & editing: SA, LC, MF, FL.

## References

Armand, S., Decoulon, G., Bonnefoy-Mazure, A., 2016. Gait analysis in children with cerebral palsy. EFORT Open Rev. 1, 448–460. 10.1302/2058-5241.1.000052

Baker, R., 2013. Measuring walking: a handbook of clinical gait analysis. Mac Keith Press, London.

Baker, R., 2006. Gait analysis methods in rehabilitation. J. NeuroEngineering Rehabil. 3, 4. 10.1186/1743-0003-3-4

Burgess-Limerick, R., Abernethy, B., Neal, R.J., 1993. Relative phase quantifies interjoint coordination. J. Biomech. 26, 91–94. 10.1016/0021-9290(93)90617-N

Byrne, J.E., Stergiou, N., Blanke, D., Houser, J.J., Kurz, M.J., Hageman, P.A., 2002. Comparison of Gait Patterns between Young and Elderly Women: An Examination of Coordination. Percept. Mot. Skills 94, 265–280. 10.2466/pms.2002.94.1.265

Carollo, J.J., Worster, K., Pan, Z., Ma, J., Chang, F., Valvano, J., 2018. Relative phase measures of intersegmental coordination describe motor control impairments in children with cerebral palsy who exhibit stiff-knee gait. Clin. Biomech. 59, 40–46. 10.1016/j.clinbiomech.2018.07.015

Chiu, S.-L., Chang, C.-C., Chou, L.-S., 2015. Inter-joint coordination of overground versus treadmill walking in young adults. Gait Posture 41, 316–318. 10.1016/j.gaitpost.2014.09.015

Chiu, S.-L., Chou, L.-S., 2012. Effect of walking speed on inter-joint coordination differs between young and elderly adults. J. Biomech. 45, 275–280. 10.1016/j.jbiomech.2011.10.028

Cockcroft, J., Louw, Q., Baker, R., 2016. Proximal placement of lateral thigh skin markers reduces soft tissue artefact during normal gait using the Conventional Gait Model. Comput. Methods Biomech. Biomed. Engin. 19, 1497–1504. 10.1080/10255842.2016.1157865

de Bruin, M., Smeulders, M.J.C., Kreulen, M., 2013. Why is joint range of motion limited in patients with cerebral palsy? J. Hand Surg. Eur. Vol. 38, 8–13. 10.1177/1753193412444401

Dixon, P.C., Loh, J.J., Michaud-Paquette, Y., Pearsall, D.J., 2017. biomechZoo: An open-source toolbox for the processing, analysis, and visualization of biomechanical movement data. Comput. Methods Programs Biomed. 140, 1–10. 10.1016/j.cmpb.2016.11.007

Dumas, R., Cheze, L., 2008. Hip and knee joints are more stabilized than driven during the stance phase of gait: An analysis of the 3D angle between joint moment and joint angular velocity. Gait Posture 28, 243–250. 10.1016/j.gaitpost.2007.12.003

Dussault-Picard, C., Cherni, Y., Dixon, P.C., 2025. Spatiotemporal characteristics of gait when walking on an uneven surface in children with cerebral palsy. Sci. Rep. 15, 4912. 10.1038/s41598-025-89280-x

Dussault-Picard, C., Cherni, Y., Ferron, A., Robert, M.T., Dixon, P.C., 2023. The effect of uneven surfaces on inter-joint coordination during walking in children with cerebral palsy. Sci. Rep. 13, 21779. 10.1038/s41598-023-49196-w

Dussault-Picard, C., Ippersiel, P., Böhm, H., Dixon, C.P., 2022. Lower-limb joint-coordination and coordination variability during gait in children with cerebral palsy. Clin. Biomech. 98, 105740. 10.1016/j.clinbiomech.2022.105740

E.A van Emmerik, R., Hamill, J., Miller, R.H., 2014. Dynamical Systems Analysis of Coordination, in: Research Methods in Biomechanics. Human Kinetics, Champaign, Illinois, pp. 291–315.

Elad, D., Barak, S., Silberg, T., Brezner, A., 2018. Sense of autonomy and daily and scholastic functioning among children with cerebral palsy. Res. Dev. Disabil. 80, 161– 169. 10.1016/j.ridd.2018.06.006

Everaert, L., Dewit, T., Huenaerts, C., Staut, L., Adams, H., Labey, L., Van Campenhout, A., Desloovere, K., 2023. Variability of gait analysis in children with Cerebral Palsy across different conditions. Gait Posture 106, S57–S58. 10.1016/j.gaitpost.2023.07.071

Fowler, E.G., Staudt, L.A., Greenberg, M.B., 2010. Lower-extremity selective voluntary motor control in patients with spastic cerebral palsy: increased distal motor impairment: Selective Voluntary Motor Control in Patients with CP. Dev. Med. Child Neurol. 52, 264–269. 10.1111/j.1469-8749.2009.03586.x

Gjesdal, B.E., Jahnsen, R., Morgan, P., Opheim, A., Mæland, S., 2020. Walking through life with cerebral palsy: reflections on daily walking by adults with cerebral palsy. Int. J. Qual. Stud. Health Well-Being 15, 1746577. 10.1080/17482631.2020.1746577

Ippersiel, P., Robbins, S.M., Dixon, P.C., 2021a. Lower-limb coordination and variability during gait: The effects of age and walking surface. Gait Posture 85, 251–257. 10.1016/j.gaitpost.2021.02.009

Ippersiel, P., Shah, V., Dixon, P.C., 2021b. The impact of outdoor walking surfaces on lower-limb coordination and variability during gait in healthy adults. Gait Posture S0966636221004859. 10.1016/j.gaitpost.2021.09.176

Konor, M.M., Morton, S., Eckerson, J.M., Grindstaff, T.L., 2012. Reliability of three measures of ankle dorsiflexion range of motion. Int. J. Sports Phys. Ther. 7, 279–287.

Lamb, P.F., Stöckl, M., 2014. On the use of continuous relative phase: Review of current approaches and outline for a new standard. Clin. Biomech. 29, 484–493. 10.1016/j.clinbiomech.2014.03.008

Leboeuf, F., Baker, R., Barré, A., Reay, J., Jones, R., Sangeux, M., 2019. The conventional gait model, an open-source implementation that reproduces the past but prepares for the future. Gait Posture 69, 235–241. 10.1016/j.gaitpost.2019.04.015

Leboeuf, F., Barre, A., Aminian, K., Sangeux, M., 2023. On the accuracy of the Conventional gait Model: Distinction between marker misplacement and soft tissue artefact errors. J. Biomech. 159, 111774. 10.1016/j.jbiomech.2023.111774

Leboeuf, F., Sangeux, M., 2023. Wand-mounted lateral markers are less prone to misplacement and soft-tissue artefacts than skin-mounted markers when using the conventional gait model. Gait Posture 100, 243–246. 10.1016/j.gaitpost.2022.12.013

Oskoui, M., Coutinho, F., Dykeman, J., Jetté, N., Pringsheim, T., 2013. An update on the prevalence of cerebral palsy: a systematic review and meta-analysis. Dev. Med. Child Neurol. 55, 509–519. 10.1111/dmcn.12080

Paiva, R., Guadagnin, E.C., Emilio De Carvalho, J., Metsavaht, L., Leporace, G., 2024. Test-retest reliability and minimal detectable change in pelvis and lower limb coordination during running assessed with modified vector coding. J. Biomech. 174, 112259. 10.1016/j.jbiomech.2024.112259

Rabbi, M.F., Pizzolato, C., Lloyd, D.G., Carty, C.P., Devaprakash, D., Diamond, L.E., 2020. Non-negative matrix factorisation is the most appropriate method for extraction of muscle synergies in walking and running. Sci. Rep. 10, 8266. 10.1038/s41598-020-65257-w

Rosenblum, M., Pikovsky, A., Kurths, J., Schäfer, C., Tass, P.A., 2001. Chapter 9 Phase synchronization: From theory to data analysis, in: Handbook of Biological Physics. Elsevier, pp. 279–321. 10.1016/S1383-8121(01)80012-9

Schmidt, A.K., van Gorp, M., van Wely, L., Ketelaar, M., Hilberink, S.R., Roebroeck, M.E., Perrin_JDecade Pip Study Groups, Tan, S.S., van Meeteren, J., van der Slot, W., Stam, H., Dallmeijer, A.J., de Groot, V., Voorman, J.M., Smits, D.W., Wintels, S.C., Reinders_JMesselink, H.A., Gorter, J.W., Verheijden, J., 2020. Autonomy in participation in cerebral palsy from childhood to adulthood. Dev. Med. Child Neurol. 62, 363–371. 10.1111/dmcn.14366

Seidler, R.D., Bernard, J.A., Burutolu, T.B., Fling, B.W., Gordon, M.T., Gwin, J.T., Kwak, Y., Lipps, D.B., 2010. Motor control and aging: links to age-related brain structural, functional, and biochemical effects. Neurosci. Biobehav. Rev. 34, 721–733. 10.1016/j.neubiorev.2009.10.005

Sint Jan, S.V., 2007. Color atlas of skeletal landmark definitions: guidelines for reproducible manual and virtual palpations. Churchill Livingstone/Elsevier, Edinburgh_J; New York.

Zeni, J.A., Richards, J.G., Higginson, J.S., 2008. Two simple methods for determining gait events during treadmill and overground walking using kinematic data. Gait Posture 27, 710–714. 10.1016/j.gaitpost.2007.07.007

